# Study of the action of lactic acid bacteria on acrylamide in food products

**DOI:** 10.1101/432898

**Authors:** Pallavi Surana, Shruti Singh, Sumathra Manokaran, A.H. Manjunatha Reddy

## Abstract

The aim of this study is to extract Lactic acid bacteria (LAB) and test its effect on starchy food products. We extracted LAB as it is GRAS (Generally Recognised As Safe) and is abundantly present in probiotics like curd. After extraction of LAB colonies and were tested for its effect on foods rich in acrylamide content. Acrylamide formation is studied by the maillard reaction. Acrylamide is found in food products which have been subjected to very high temperatures. The application of LAB to remove many mutagens, by using the binding mechanism was assessed for acrylamide. When this acrylamide degrades it produces toxic gases like ammonia, hydrogen and carbon monoxide. Asparagine is an important precursor to acrylamide formation. So in foods containing free asparagine group with reducing sugars will form acrylamide. We used liquid chromatography to determine the presence of acrylamide in the food product. Acrylamide content in the body is responsible for various mutations at the genomic level thereby, leading to cancers like colon, breast. It also leads to neural damages and is directly linked to Alzhimers disease. This study concludes the beneficial effects of consuming probiotics and suggests a way to reduce the acrylamide formation in food products.

## Introduction

Acrylamide is a carcinogenic substance usually formed when a sugar and amino acid react at extremely high temperatures. [1] It undergoes a chemical reaction called the Malliard reaction with a string focus on asparagine, as studies suggest. [2] This reaction is responsible for the taste, flavour and aroma of the food. This chemical is found in bread, certain starchy foods like potato which is deep fried, biscuits baked at high temperatures. The acrylamide content was discovered in certain types of food cooked at high temperatures in April 2002 by the Swedish National Food Authority. [3] It was a ground-breaking discovery as now it was certain that cooking food at very high temperatures does not enhance the taste only but can be lethal, too.

The first step in formation of acrylamide is the formation of a Schiff base. This base can take two processes from here, it can either hydrolyze to form 3-aminopropionamide that can further degrade via the elimination of ammonia to form acrylamide when heated. Alternatively (as shown in Fig 1) the decarboxylated Schiff base can decompose directly to form acrylamide via elimination of the imine. Temperature is an important factor in the formation and elimination of acrylamide formation or not. It usually forms in foods cooked at temperatures beyond 120 degree Celsius. [2] Hence its known that asparagine is often considered as a precursor for acrylamide synthesis.

**Fig.1.**
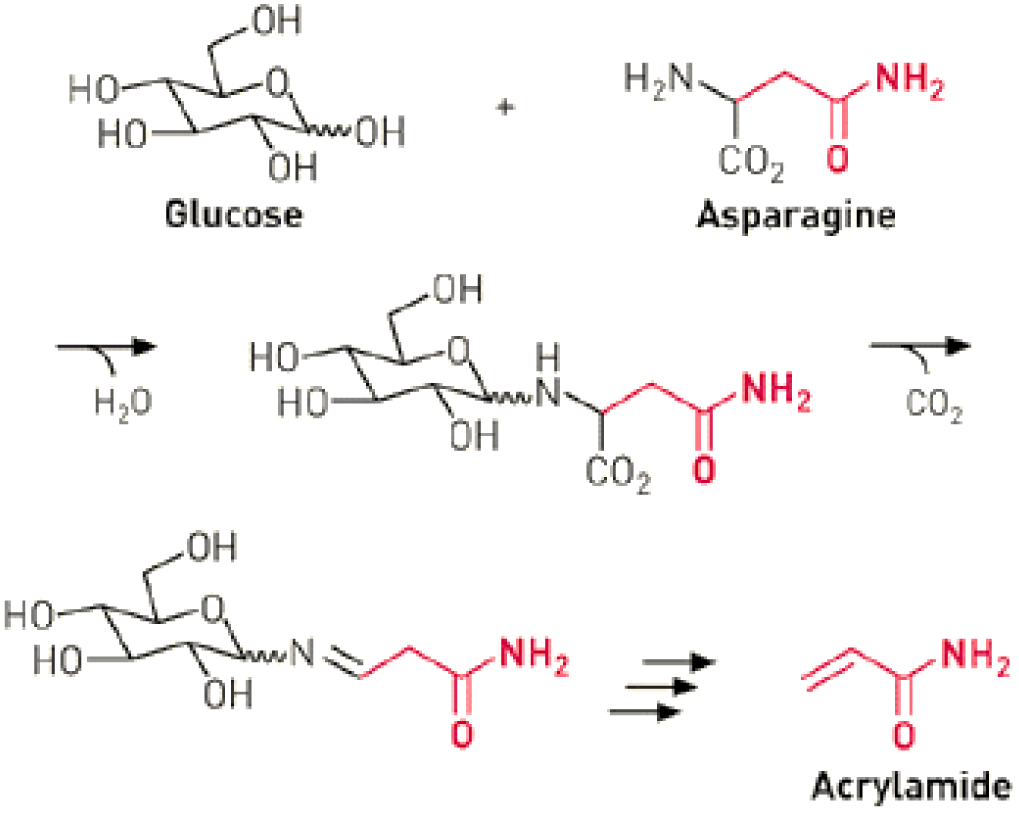
Mechanism of acrylamide formation - The pyrolysis of asparagine with glucose [4]

Zeracryl a company worked on a patented technology which proved to use probiotics like LAB (lactic acid bacteria). This LAB is abundantly found in curd and various other milk products, curd being the most enriched one. The method spoke about how food-grade LAB can reduce the concentrations of reducing sugars like glucose on the surface of foods. This would thereby, reduce the acrylamide formation at high temperatures. The company has successfully proven that the method is 90 % effective in use. [5]

This study focuses on extraction of LAB from curds and the testing its effects on acrylamide containing foods like starch rich foods cooked at high temperatures. This importance of this study lies in the fact that dietary acrylamide is known to cause cancers and Alzhimers disease. As known from ages that acrylamide causes muscle weakness and decreased muscle coordination. It has also been found that it causes degradation of nerve cells and thereby might lead to development of neurodegenerative disorders. [6]

## Methodology

### Isolation of LAB from curd

A number of sources can be used to extract LAB. This study focuses on extraction of LAB from curd. LAB is a gram positive bacterium that grows well in aerobic and anaerobic conditions. [7]

The first step is to suspend the curd samples in diluted in sterile saline solution. Serial dilutions are made for up to 6 times. This is done by taking 1ml of sample and adding saline of the remaining to make up to 10 ml. This is then inoculated on a media. There are various types of media that can be used like MRS agar, Rogosa Agar. The preferred one is the Mann Rogosa Sharpe (MRS) Agar. [8] Then the colonies were stored in the agar medium along with glycerol at −20 degrees Celsius. Later the bacteria was extracted from the agar and purified.

### Action of Lactic Acid bacteria on acrylamide in food

The process followed next was to check how the LAB would work on the acrylamide containing foods. A small quantity of the sample was loaded in a a mixing solution. The solution is usually made up of KOH and ethyl alcohol. Heat is them applied to this to hydrolyse the sample. The sample can also be homogenised with distilled water.

The next step was to centrifuge the sample and maintain a pH of about 6.5 to 6.8. This pH is usually achieved by adding acids like HCl. This was done to the aqueous phase. The next step is to precipitate the proteins and carbohydrate molecules, for this we use the Carrez solutions I and II in 1:1 ratio. [9,10] The Carrez Reagents are used for the analysis of small molecules such as carbohydrates, alcohols, aldehydes and organic acids. [11]

This mixture was shaken vigorously and then the sample was centrifuged after which we take 10 ml of the filtered sample. This filtered sample was shaken for about 30-35 minutes in a mechanical shaker. This gives a homogenised solution. The derivitisation reaction is carried out at the suitable room temperature. We add 100 microlitres of 6M HCl, 2μL of acrylamide-d3. The derivatization reaction was maintained for 30 to 35 minutes at appropriate room temperature. The solution is the neutralised with a base preferably, KOH. The pH of the solution is further mainatined to 9.0 by using NaHCO3/K2CO3 (3:1, w/w). The DLLME procedure was applied to extract of xanthyl-acrylamide. The extracting solvent used is tetracholoroethylene and is extracted with acetone as the major solvent which is the disperser solvent. The dispersed particles were sediment, then the GC-MS analysis was done. [9]

Another method widely used is HPLC (high pressure liquid chromatography). Here after the addition of Carrez I and II solutions, hexane was added. This hexane extracts the long chain fatty acids and is very effective. The upper layer of hexane is removed after vigorous mixing. This layer contains the unwanted fatty acids. The solution is filtered and added to the HPLC column using a thin needle or a syringe.

### Effect of pH, water and fermentation on the acrylamide content

It has been observed in research studies that the pH affects the acrylamide content. Any reduction of pH, increases acidity and thereby increases the possibility of formation of Maillard reaction associates. This is supplemented by the formation of the carcinogen, acrylamide. Effect of water also plays a role in acrylamide formation.[12] When the food is cooked at very high temperatures, it tends to lose its moisture content. The moisture content between 0.4 to 0.7 is the range where there is acrylamide activity observed. Fermentation similarly for long hours can favour acrylamide formation. It is observed that acrylamide removal is highest from potato with water content of 0.95 which is 20%. [13]

## Discussion

In this review we tried to integrate the methodology of extracting LAB from a milk product like curd. The acrylamide produced also depends on various factors like pH, temperature, moisture content. The GC-MS and HPLC methods quantify the amount of acrylamide content. The food items with high concentration of the acrylamide have been reduced its acrylamide concentration due to introducing the lactic acid bacteria.

